# Photoredox-catalyzed decarboxylative *C*-terminal differentiation for bulk and single molecule proteomics

**DOI:** 10.1101/2021.07.08.451692

**Authors:** Le Zhang, Brendan M. Floyd, Maheshwerreddy Chilamari, James Mapes, Jagannath Swaminathan, Steven Bloom, Edward M. Marcotte, Eric V. Anslyn

## Abstract

Methods for the selective labeling of biogenic functional groups on peptides are being developed and used in the workflow of both current and emerging proteomics technologies, such as single-molecule fluorosequencing. To achieve successful labeling with any one method requires that the peptide fragments contain the functional group for which the labeling chemistry is designed. In practice, only two functional groups are present on every peptide fragment regardless of the protein cleavage site, namely, an *N*-terminal amine and a *C*-terminal carboxylic acid. Developing a global-labeling technology, therefore, requires one to specifically target the *N*- and/or *C*-terminus of peptides. In this work, we showcase the first successful application of photocatalyzed *C*-terminal decarboxylative-alkylation for peptide mass-spectrometry and single molecule protein sequencing, that can be broadly applied to any proteome. We demonstrate that peptides in complex mixtures generated from enzymatic digests from bovine serum albumin, as well as protein mixtures from yeast and human cell extracts, can be site-specifically labeled at their *C*-terminal residue with a Michael acceptor. Using two distinct analytical approaches, we characterize *C*-terminal labeling efficiencies of greater than 50% across complete proteomes and document the proclivity of various *C*-terminal amino acid residues for decarboxylative-labeling, showing histidine and tryptophan to be the most disfavored. Finally, we combine *C*-terminal decarboxylative labeling with an orthogonal carboxylic acid labeling technology in tandem, to establish a new platform for fluorosequencing.

## INTRODUCTION

Progress in protein mass spectrometry, along with the development of single-molecule protein sequencing technologies, and other advancements in protein detection and quantification, has yielded several new opportunities for proteomics^1,2^. Examples include: (1) the discovery of novel microproteins^3^, (2) the deployment of single-cell proteomics^4,5^, and (3) the discovery of new diagnostic tools for the clinic^6^. The vast majority of existing applications take advantage of covalent labeling technologies for peptides and proteins, many of which are designed to target specific functional groups found on biogenic amino acid (AA) residues. While extant conjugation methods have certainly opened the door to new possibilities for protein mass spectrometry, still more methods are needed to mine this nascent area of research. For mass spectrometry, the need for innovative cross-linking strategies to study protein-protein interactions and methods to label peptides with isotopic residues for quantitation and multiplexing are at the top of the list.^7,8^ The development of single-molecule protein sequencing methods has also led to a pronounced need for labeling chemistries that can discriminate between specific amino acid residues found in peptides,^12,13^ particularly those that share a common functional group (e.g., the amines of lysine and *N*-terminal residues, and the carboxylic acids of aspartic acid, glutamic acid, and the *C*-terminal residue)^2^. One way to address these topics would be to develop, and/or to make use of, labeling technologies that specially target the *C*- or *N*- terminus of peptides. These termini are: (1) naturally present in almost every peptide, (2) often removed from the active binding site of the peptide, meaning that labeling these positions would be less likely to interfere with the endogenous activity of the peptide, and (3) generally accessible for modification, free from any steric or conformational constraints^9–11^.

A number of techniques have been developed for the selective and efficient labeling of the *N*-terminal amino group of peptides and proteins (as opposed to the amine of lysine residues)^14,15^. Corresponding methods for attaching a synthetic reporter to the *C*-terminus of peptides and proteins that also keep alternate side chain acidic residues (i.e., aspartate and glutamate) pristine are few and seldom well optimized^16–18^. Such discrimination is intrinsically challenging as the reactivities of carboxylic acid side chains and *C*-terminal carboxylates are very similar, and at least for full-length proteins, the acidic side chains are considerably more abundant than free *C*-termini. Hence, there are several innate chemical challenges to specifically derivatize protein or peptide *C*-termini, although the benefits from doing so are significant, especially for proteomics (e.g., isotopic quantitation, multiplexing, and sequencing).

The selective labeling of *C*-terminal carboxylic acids of peptides in complex peptide mixtures for shotgun proteomics has been reported in the literature^19^. One such method involves oxazolone formation via nucleophilic addition of *C*-terminal carboxylic acids to acetic anhydride^20,21^. Another method involves enzymatic coupling with carboxypeptidase, wherein the enzyme ligates a nucleophilic molecule to the *C*-terminal carboxylic acid at high pH^17^. However, in our experience, these methods lack the necessary *C*-terminal selectivity and mild reaction conditions that are required for applications to advanced proteomics. In our hands, we observed an indiscriminate cross-reaction of acetic anhydride with any number of peptide nucleophilic amino acid side chains, which blocked many ionizable groups, thus deterring subsequent conjugation reactions and hindering analysis by conventional mass spectrometry. Similarly, the enzymatic method was synthetically laborious to implement—required time-consuming preparations prior to labeling, including esterification and lyophilization procedures—and the dramatic change in pH proved unsuitable for some peptides and proteins that were more pH-sensitive.

Recently, new methods have been described that offer the potential for faster, more generalized *C*-terminal modification via photoredox catalyzed decarboxylation of *C*-terminal carboxylic acids^22–24^. These methods have been described for small peptides (most less than 10 amino acids), but offer a starting point for applications to proteins, and the field of proteomics in general. Herein, we detail our studies to optimize and to assess the feasibility of photocatalyzed *C*-terminal labeling for bottom-up proteomics, simultaneously modifying and characterizing entire libraries of peptides from complex protein digests. We extensively describe the methods, instrumentation, and expectations for success when applying decarboxylative-alkylation for bottom-up proteomics, such that it can be widely adopted by others.

## RESULTS AND DISCUSSION

### Optimizing the conditions for angiotensin coupling

We first sought to optimize the reaction conditions starting from the procedure reported by Bloom *et al*^22^. The photoredox reactions are carried out in a commercially available instrument—Lumidox II system (Analytical Sales and Services, New Jersey)—fitted with an active cooling base (Analytical Sales and Services, NJ), and 445 nm Blue LEDs. A table fan was used to keep the reaction block cooled to room temperature for the duration of the experiment (6 hr.). (Note: Lumidox II allows for up to 96 photoredox reactions to be completed in parallel and is an optimal platform for small-scale reaction optimization and performance.)^25^ A photograph of the setup is shown in **Figure 1A**. The small peptide angiotensin-II (seq: H_2_N-DRVYIHPF-CO_2_H) and the Michael acceptor 3-methylene-2-norbornanone (herein termed NB for short) were used to screen experimental conditions for efficient conversion (reaction scheme shown in **Figure 1B**). The effect of aqueous buffers with various co-solvents, amount of catalyst loading, equivalents of Michael acceptor, and angiotensin concentration were all varied to optimize the reaction (summarized in Supplementary Tables ST1-ST4). A representative LC trace is shown in **Figure 1C**, which highlights the conversion efficiency to the angiotensin-NB adduct by comparison to the peaks corresponding to endogenous (unmodified) angiotensin and decarboxylated angiotensin peptide. The mass chromatogram of the corresponding LC trace is provided in **Supplementary Figure SF1**. The experiments with cesium formate buffer (pH 3.5) and DMSO (5%, 10%, 20% v/v) resulted in high conversion. Considering that greater amounts of DMSO may diminish signals of modified and unmodified peptides, we elected to use only 5% DMSO (v:v) for further reactions.

**Figure 1.**
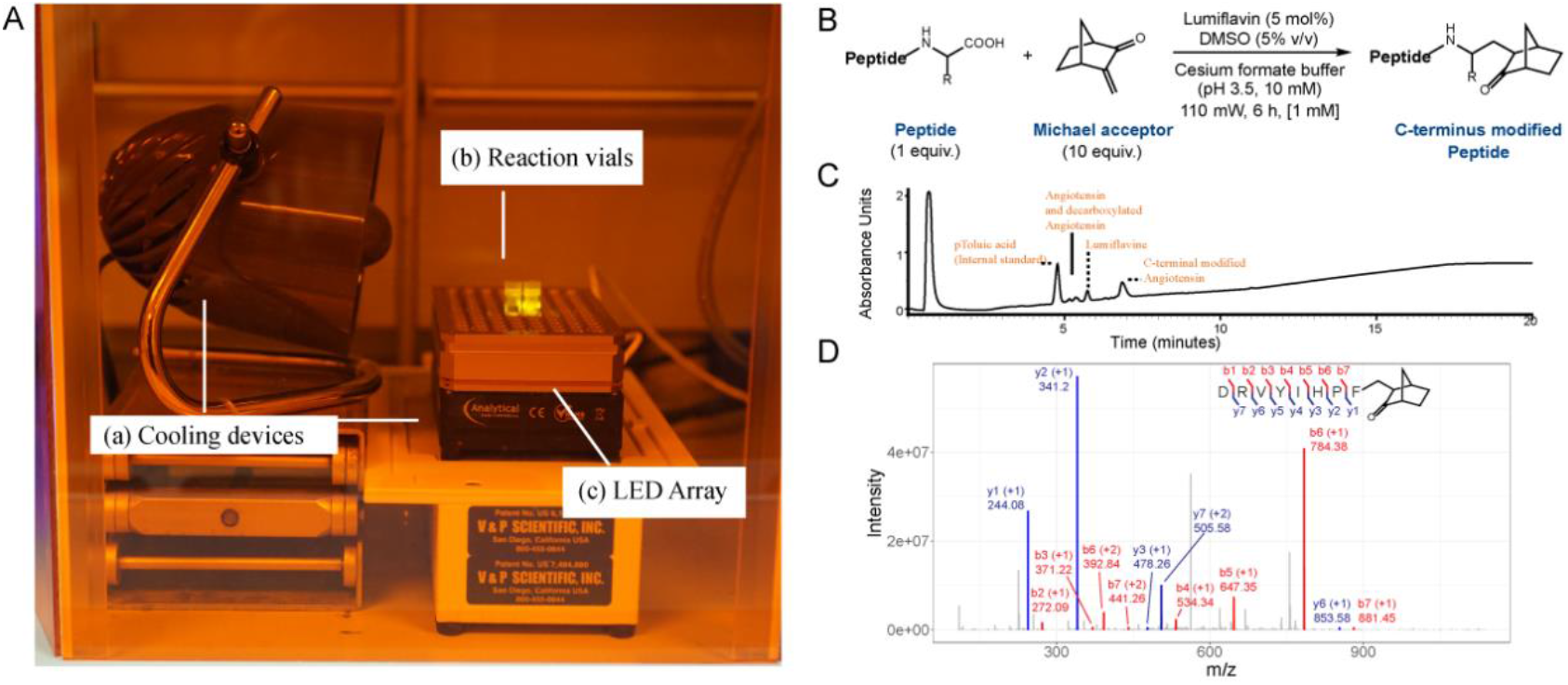
Demonstrating the selective *C*-terminal carboxylic acid labeling of angiotensin. (A) Reaction setup for performing the photoredox chemistry. The three main components as indicated in the picture are - (a) a tabletop fan and an active cooling unit, (b) a glass vial containing the reaction mixture, including the photocatalyst lumiflavin, peptide, and Michael acceptor NB in acidic buffer, and (c) the Lumidox Gen II LED array with aluminum block (Analytical Sales and Services, NJ, USA). (B) Reaction scheme illustrating derivatization of the terminal carboxylic acid with NB. (C) UPLC-Chromatogram of the reaction products. (D) Tandem (MS/MS) mass spectrum confirming modification of the terminal carboxylic acid

Higher conversions (81%) could be achieved by increasing the catalyst loading to 10 mol % and using 10 equiv. of Michael acceptor (1 mM concentration). Finally, we optimized the reaction time and lamp power. We determined that the reaction performed the best with 110 mW/well and 6 hours, giving an average conversion of ~60%. High resolution MS-MS results of the angiotensin-NB adduct indicated that the *C*-terminus of angiotensin was selectively modified (**Figure 1D**). The optimized conditions were reproducible at two independent locations (in setups at the University of Kansas and at UT-Austin).

### Determining amino acid biases in the reaction chemistry

After initial optimization of the photoredox reaction, we suspected that there could be innate biases with respect to the *C*-terminal amino acids, with some *C*-terminal amino acids being more susceptible to decarboxylative-alkylation. Based on the mechanism proposed in the literature^22^, we rationalized that this bias could result from the stability of distinct α-amino radicals generated from different *C*-terminal residues (affecting both the rate of decarboxylation and reactivity of the α-amino radical toward conjugate addition), as well as inherent differences in oxidation potentials between the various *C*-terminal residues. Thus, we investigated the variance in the yields of twenty-one short peptides, 20 of which have a unique biogenic amino acid at their *C*-terminus and 1 of which has its *C*-terminal acid replaced by an amide to serve as a negative control. The conversions are summarized in **Table 1**.

**Table 1:**
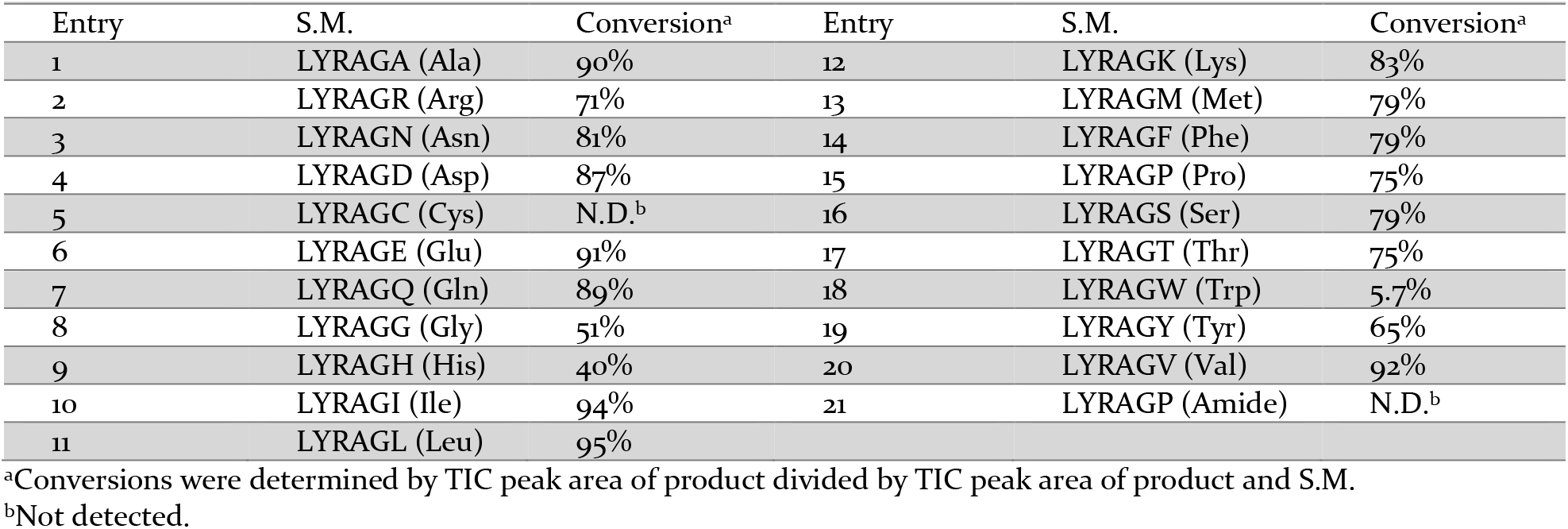
Determination of conversions of peptides with varying *C*-terminal amino acids.

The data reveals that 13 of the 20 peptides gave greater than 75% conversion upon reaction with NB. The reaction of the *C*-terminal Cys-containing peptide with NB did not furnish the *C*-terminal modified product. Competitive thiol-conjugate addition was observed instead. Therefore, one will need to protect thiol groups to proceed with this *C*-terminal diversification strategy, as described below using iodoacetamide. It is worth noting that a tryptophan *C*-terminal peptide gave a much lower conversion of modified product (~6%), one reason being the fast and reversible electron transfer between the electron-rich indole and the lumiflavin photocatalyst. Protonation of the imidazole ring of the *C*-terminal histidine peptide at pH 3.5 could deter single-electron transfer oxidation and might also diminish the nucleophilicity of the α-amino radical, and these issues could explain the lower conversions observed in this case (40%). As anticipated, a peptide lacking a *C*-terminal carboxylic acid gave no product.

### Modifications occur efficiently on *C*-terminal carboxylic acids in heterogeneous peptide digests

To gain a better understanding of the applicability of the *C*-terminal differentiation reaction for proteomics experiments, we performed the reaction on heterogenous peptide samples and probed the extent of the modifications using tandem mass spectrometry (LC-MS/MS). We digested the protein bovine serum albumin (BSA), proteins from yeast lysate, and human cell lysate with the endoproteases trypsin and GluC. In most cases trypsin cleaves after lysine and arginine (K/R), and GluC cleaves after aspartate and glutamate (D/E). We then split these samples into control and experimental samples where the only difference between the two samples was the presence or absence of the catalyst lumiflavin. After desalting and C-18 tip exchange, we analyzed the peptides using LC-MS/MS and searched for the NB modification in the analysis software as a *C*-terminal mass addition of 78.083 Daltons (**Fig. 2a**). For each sample we were able to estimate the *C*-terminal labeling efficiency as a ratio of peptide spectral matches (PSMs) with a *C*-terminal NB and the total PSMs observed in a sample. In the trypsin digested samples, we observed an average labeling efficiency for each of our experimental digests ranging from 52% for the BSA sample to 76% for the human lysate (**Fig 2b**). Labeling efficiencies in GluC digested samples ranged from 65% in the human lysate sample to 89% in the yeast lysate sample (**Fig 2c**). The difference in labeling efficiencies between these two sets of experiments may arise from the different terminal amino acids generated by the two proteases, as suggested by the analysis of **Table 1**. However, the high extent of labeling should support most basic proteomics experiments.

**Figure 2.**
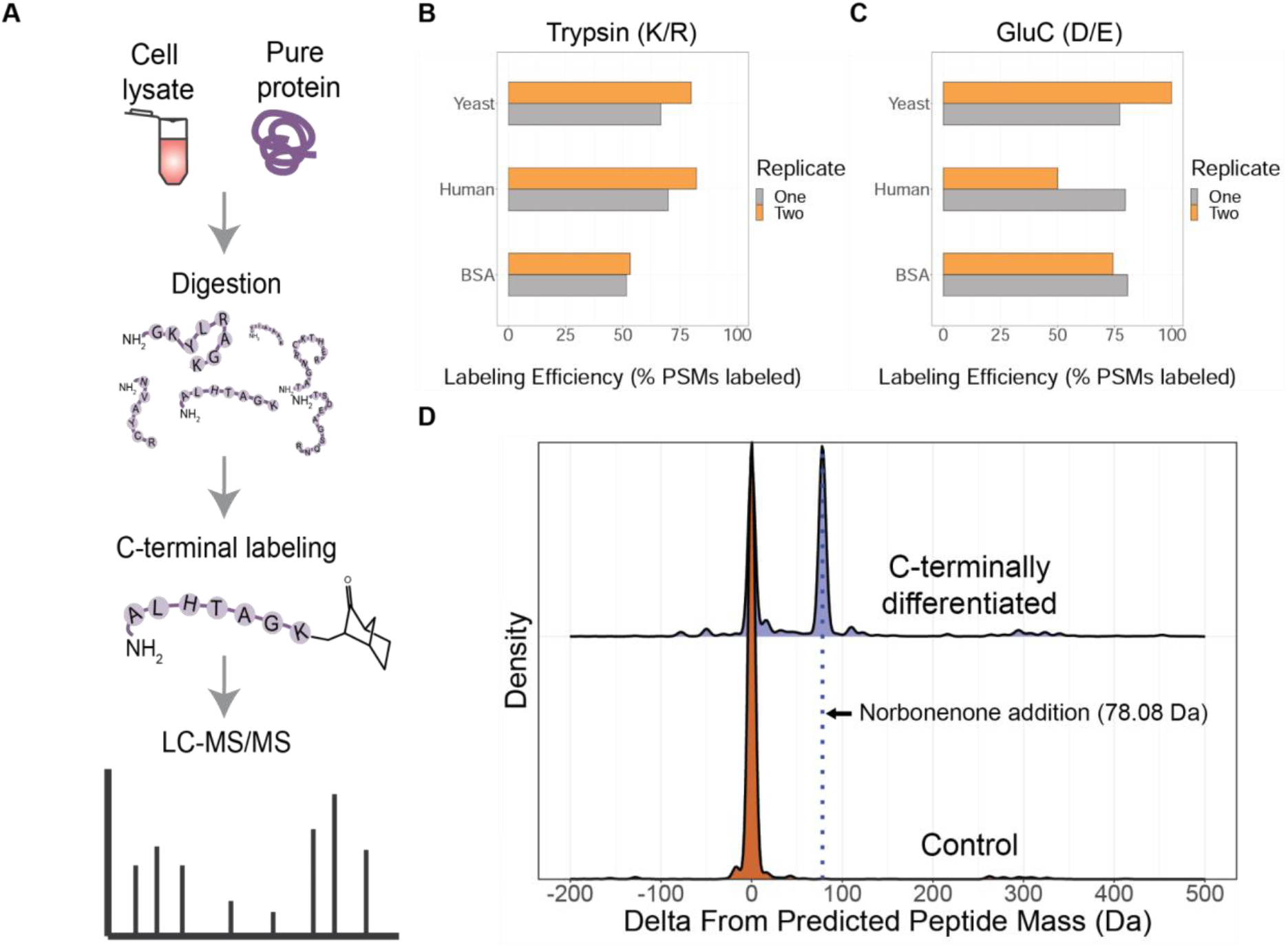
C-terminal carboxylic acid modifications occur with high efficiency on proteomic digests. (A) Workflow of the proteomics experiment with C-terminal labeling. C-terminal differentiation reaction labeling efficiency on (B) trypsin and (C) GluC digested yeast, human, and BSA protein samples. Experiments done in duplicate. (D) Delta mass values from predicted peptide mass for the C-terminally differentiated sample (purple) and the control sample (orange). Blue line corresponds to the NB mass addition of 78.08 Da.

Interestingly, when comparing the total peptide abundances between our negative control and experimental samples we consistently saw, on average, a 76% reduction in total high confidence PSMs in our experimental samples (**Supplementary Figure SF2**). We considered multiple hypotheses as to the cause of this decrease in peptide identifications. The first of these was that there were side reactions occurring in our experimental sample resulting in unexpected modifications on peptides that would lower the confidence of identification. The second was that the proteomics software could not localize the *C*-terminal modification resulting in a decrease in high confidence PSMs. The third was that 445 nm irradiation in our experimental setup degraded some of our peptides in the presence of lumiflavin. To test the first of these hypotheses we sought to identify what, if any, side reactions might have occurred during the reaction process with our previously collected shotgun proteomics data. Thus, we reanalyzed the mass spectra using the proteomics software MSFragger^26^, performing an open search (see Methods) to identify all chemical modifications — without specifying them in advance — that resulted in a change in mass from the predicted, unmodified peptide masses. Two distinct peaks were observed. One was centered on 0 Da corresponding to the unmodified peptide, and the other was centered at 78.08 Da closely corresponding to the expected mass addition of NB. In the control sample, we observed only a single large peak centered on 0 Da, as expected (**Fig 2d**). This analysis confirmed that the primary reaction observed was the expected *C*-terminal addition of NB and that our modification reaction only occurred in the presence of lumiflavin photocatalyst. Based on the MSFragger analysis, we could thus rule out the presence of appreciable side reactions leading to altered modification sites. Notably, even in the MSFragger open searches we still observed a 73% average decrease in peptide identifications. These analyses thus suggest that some degradation due to irradiation in the presence of lumiflavin may cause a reduction in peptide abundances. Overcoming this limitation may call for the use of an alternative flavin-photocatalyst, narrowed irradiation window, and other slight changes to the experimental protocol, which will serve as a starting point for follow-up studies. In spite of the reduced yield, the reaction proved to be highly selective when labeling the *C*-termini of peptides in a heterogenous biological sample.

### Minimum *C*-terminal amino acid bias occurs on peptides from biological samples

We next sought to determine if the *C*-terminal biases we observed with synthetic peptides in **Table 1** were also reflected in the biological samples. Any use of the decarboxylative-alkylation chemistry on intact proteins and/or peptides or proteins whose terminal amino acids vary will require minimal bias in the reaction chemistry for accurate quantification, particularly in fields such as *C*-terminomics^19^. To evaluate the *C*-terminal bias in complex mixtures, we again digested BSA, yeast, and human protein extract. To generate non-uniform *C*-terminal amino acids we used three proteases that cut specified amino acids downstream of the *N*-terminus: LysN (K), AspN (D/E), and LysargiNase (K/R). Using these proteases, we were able to generate a mostly random population of *C*-terminal amino acids and estimate the labeling efficiency of the *C*-terminal differentiation reaction. For most amino acids the median labeling efficiency was between 75-100% (as seen in **Figure 3**). In certain cases, such as with histidine and methionine, the labeling efficiency was much lower than expected based on the other amino acids. For methionine, this is likely an artifact of low sampling rather than the actual labeling bias as the earlier experiments with the purified peptide showed an elevated labeling efficiency (**Table 1**). The histidine result corresponds with that observed in the purified peptide and was discussed above. It should be noted that tryptophan was only observed once across all of the experiments and so was left out of this analysis, but it would have been expected to not be efficiently labelled based on **Table 1**. The high labeling efficiency across most of the *C*-termini demonstrates the robustness of the differentiation reaction. The largely unbiased nature of the reaction chemistry across peptides makes it highly attractive to methods such as *C*-terminal peptide enrichment strategies and single molecule protein sequencing.

**Figure 3.**
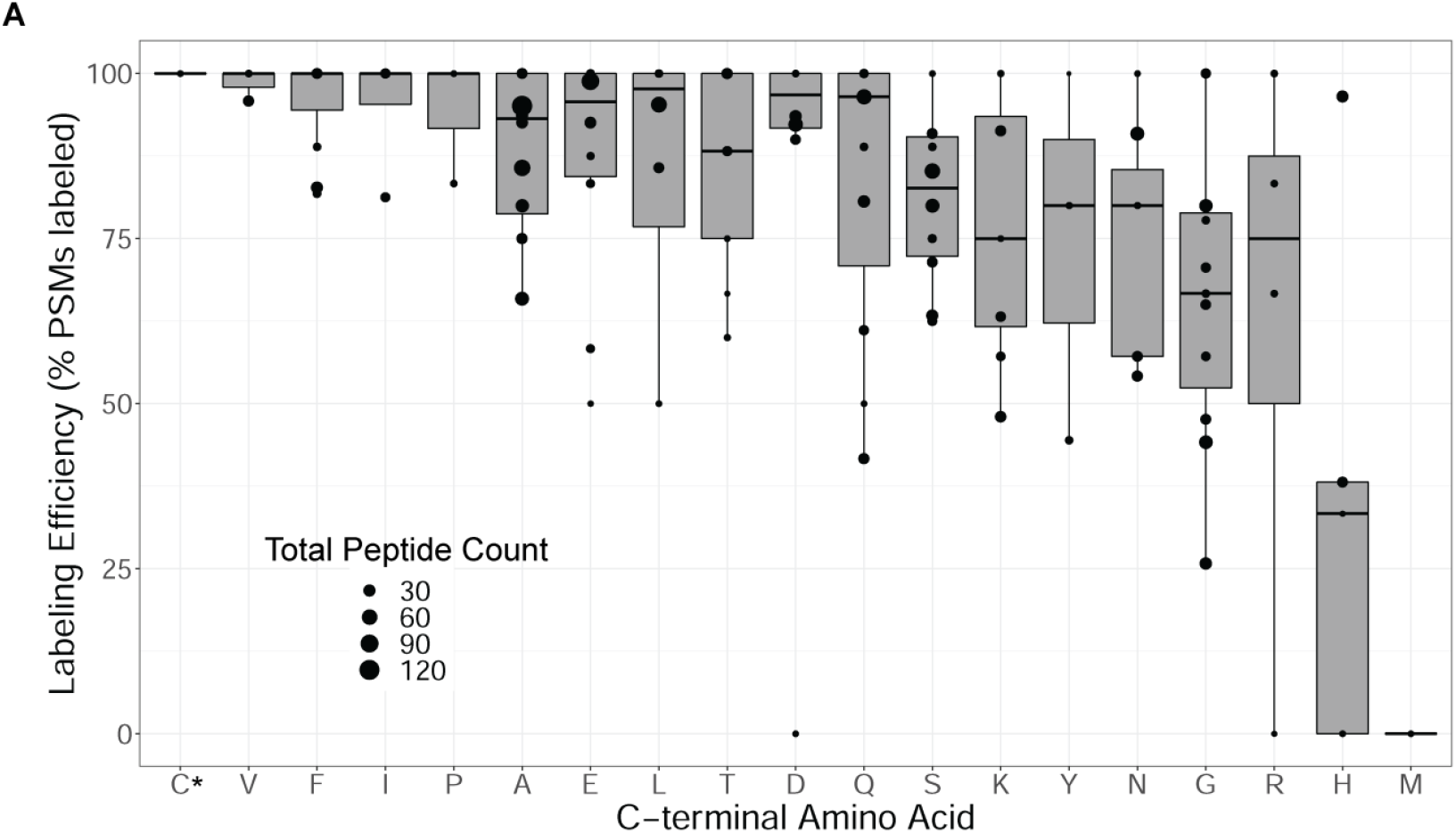
*C*-terminal labeling exhibits only moderate biases across peptides with varying *C*-terminal amino acid residues. The plot summarizes analyses of peptides generated from 15 independent experiments from digesting protein samples (BSA, yeast, and human cell protein extracts) using *N*-terminal cleaving proteases (AspN, lysN, and LysargiNase) that expose varying peptide *C*-terminal amino acids. Peptides were identified by tandem mass-spectrometry and the frequency of each *C*-terminal modified amino acid to its overall observed frequency was calculated to determine the labeling efficiency. Labelling efficiencies were determined for each of the 15 experiments, indicated as separate data points. C* - Cysteine has been alkylated in advance using iodoacetamide.

### Adapting the method for single molecule protein sequencing

Our group recently developed a new proteomics technology, termed *fluorosequencing*, for identifying peptides in a complex mixture at a single molecule resolution^9^. In this method, a researcher selectively labels amino acids on peptides with fluorophores, immobilizes the peptides on a glass slide, subjects them to rounds of Edman degradation chemistry, and monitors the fluorescence of individual peptides after loss of each consecutive amino acid per Edman cycle. The Edman cycle coinciding with the loss in fluorescent intensity (occurring due to the removal of the fluorescently labeled amino acid) indicates the position of the labeled amino acid in the sequence. This pattern, termed as a *fluorosequence*, is mapped to a reference database to identify the likely source peptide and proteins. As can be observed in the workflow, selectively labeling amino acids with fluorophores and immobilizing the peptides to a solid imaging surface are two critical steps in the process. The selective labeling of the *C*-terminal carboxylic acid not only provides a unique attachment point for immobilization, but its exclusivity allows for subsequent labeling of internal acidic residues using alternate technologies, such as amide coupling, which can enhance the fluorosequencing method when used in tandem. We implemented the peptide labeling process for angiotensin (illustrated in **Figure 4A**) by first conjugating the *C*-terminal carboxylic acid with a synthesized NB-PEG4-alkyne linker in solution using the above described and optimized photoredox chemistry, then capturing the peptide on solid support via’ its *N*-terminal amine using a pyridine carboxaldehyde reagent (PCA) (procedure adapted from^15^), followed by a two-step labeling process to attach a fluorophore on the glutamic acid. The two-step coupling reaction involves an amide coupling using 3-azidopropylamine followed by installing an Atto647N-PEG4-DBCO using standard copper-free click chemistry. Following the labeling step and washing away the free uncoupled fluorophores from the solid support, we released and deprotected the peptide scarlessly from the support. After a serial dilution through four orders of magnitude, we immobilized the liberated and fluorescently labeled peptide on an azide-functionalized glass surface using copper-catalyzed click chemistry (see methods). Fluorosequencing this peptide (seen in the blue bar in **Figure 4B**) reveals the position of the glutamic acid residue on individual molecules of angiotensin, as seen by the preferential removal of the fluorophore after the 1st Edman cycle. As suitable negative controls, we also fluorosequenced: (a) angiotensin but without the alkyne linker to control for non-specific attachment of the labeled peptide on the azide slides, and (b) a negative control peptide of sequence H_2_N-AGAGANGSNFGAN-(CO)NH_2_, with a amidated *C*-terminal carboxylic acid to serve as a control for excess fluorophores carried along through the reaction. The 20-fold increase in the counts of fluorescence loss after the 1st Edman cycle for angiotensin when compared with the controls confirms the utility of the selective *C*-terminal decarboxylative-alkylation chemistry for enabling single molecule sequencing of internal carboxylic acid-containing amino acid residues.

**Figure 4.**
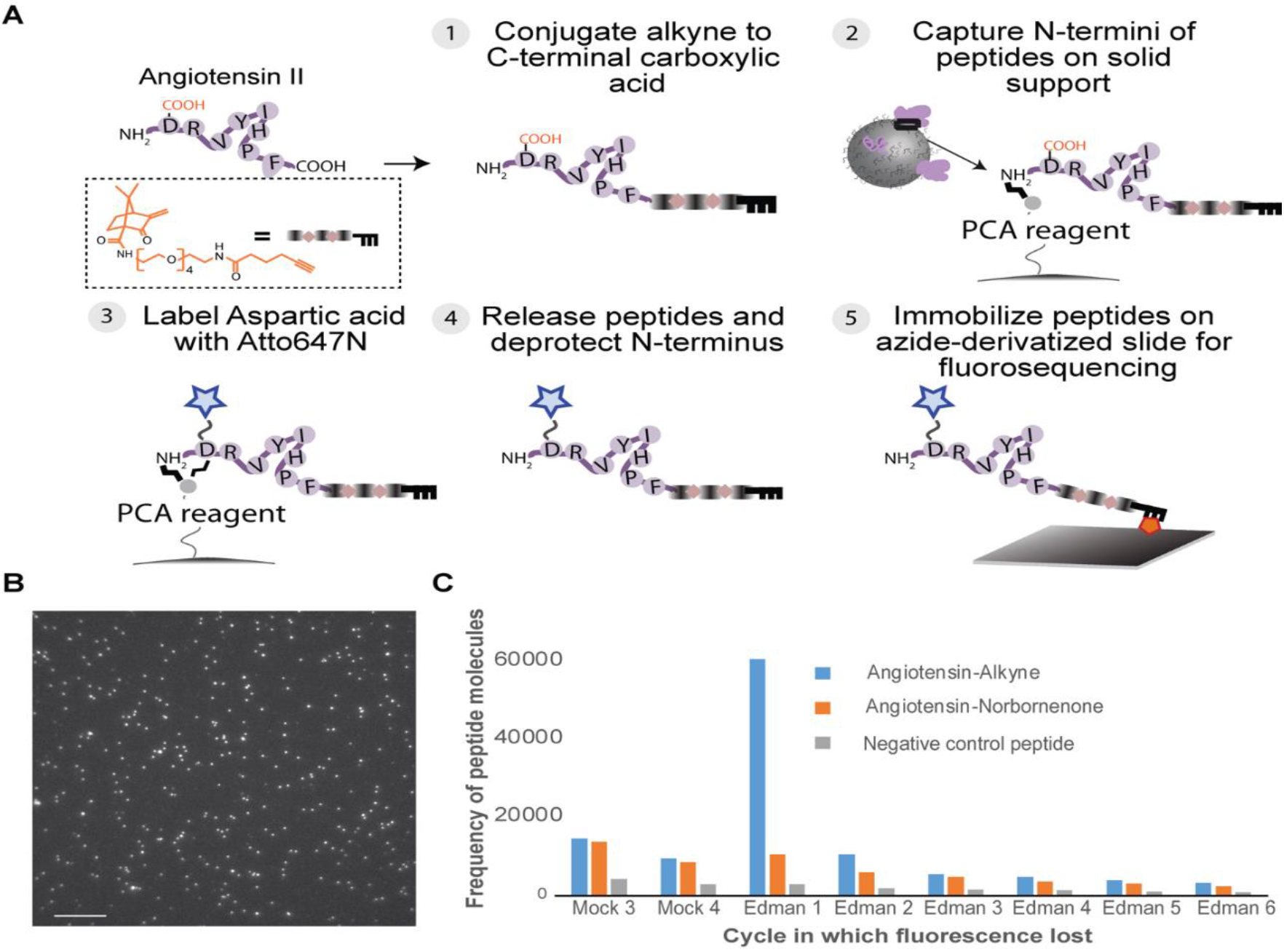
Application of *C*-terminal differentiation to immobilize peptides for fluorosequencing, enabling single molecule determination of the sequence position of fluorophore-derivatized internal acidic amino acids. The *C*-terminal carboxylic acid of angiotensin was labeled with NB-PEG4-alkyne and its internal aspartic acid was labeled with fluorophore Atto647N prior to single molecule peptide sequencing. (A) The steps involved in the process are illustrated. (B) A representative image of the numerous fluorescently labeled single peptide molecules of angiotensin labeled with Atto647N and immobilized on an azide functionalized glass slide. (C) The bar chart shows the result of the fluorosequencing experiment performed on this labeled and immobilized angiotensin molecules, with (angiotensin - alkyne, blue bars) and without alkyne (red bars) and negative control (peptide - AGAGANGSNFGAN-(CO)NH_2_, gray bar) which is incapable of *C*-terminal functionalization as well as labeling.The peptides were exposed to four cycles of “mock” Edman degradation (performing the full set of treatments but omitting phenyl isothiocyanate; indicated by cycle names beginning with “M”) followed by six cycles of Edman degradation (indicated by cycle names beginning with “E”). The position of the Edman cycle that corresponds to the maximal loss of peptides indicates the position of the fluorescently labeled amino acid. The results demonstrate the successful implementation of the discriminative carboxylic acid labeling and its use in single molecule protein identification technology. The scale bar denotes 10 μm.

## CONCLUSIONS

Photoredox catalyzed decarboxylative-alkylation has been reported to label the C-terminal carboxylic acid residue of peptides with high efficiency, but the vast applications of this methodology are untapped. With the goal of applying this approach to the field of bottom-up proteomics and in other emerging single molecule proteomics technologies, we first optimized the photocatalyzed procedure using endogenous angiotensin as our target substrate. We then tested this method with 21 synthetic peptides, each having a chemically unique *C*-terminal residue, and compared the labeling efficiencies. Following this, we applied the decarboxylation protocol to biological samples and estimated the labeling efficiency in these complex mixtures. A search of possible alternative modifications verified that there were no prominent side reactions. We then used a diverse set of proteases to generate and to characterize a mostly random pool of *C*-terminal peptides in a biological sample, in terms of labeling efficiency and bias. These results both showed minimal amino acid bias in complex solutions and corresponded closely to the labeling biases observed with purified peptides. Finally, we demonstrated the utility of decarboxylative-alkylation for single molecule proteomics (namely, fluorosequencing) by using this chemistry to append an alkyne-containing linker to the *C*-terminus of angiotensin II, and then using orthogonal chemistries to couple and to sequence the peptide on-slides using TIRF microscopy. We anticipate that this differentiation reaction will be broadly useful for proteomics applications.

## METHODS

### Materials used

The main materials purchased were angiotensin (Sigma, Cat #05-23-0101), 3-methylene-2-norbornanone (Sigma; Cat# M46055), lumiflavin (Cayman Chemical, Cat# 20645) and Cesium formate (Alfa Aesar, Cat #B2132714). All synthetic peptides were synthesized by Genscript (NJ, USA). Intact protein and trypsin/LysC digested peptides for BSA (Cat# 88341, Thermo), yeast (*Saccharomyces cerevisiae*) cell protein extract (Cat# V7341, V7461), and human K562 cell protein extract (Cat# V6951, V6941) were purchased from Promega Inc (WI, USA). NB-PEG4-Alkyne linker was custom synthesized and purified by Comminex Inc (Budapest, Hungary) based on the procedure described in Bloom *et al*. [29359756]. Proteases Pierce trypsin (Cat #90058), Pierce GluC (Cat #90054) and Pierce LysN (Cat#90300) were purchased from Thermo Scientific. LysArginase (Cat #EMS0008) was purchased from Millipore Sigma (WI, USA). AspN (Cat#) was purchased from Promega Inc (Cat #V1621). 2,2,2-trifluoroethanol (Cat #T63002) and tris(2-carboxyethyl)phosphine (Cat #77720) were purchased from Sigma Aldrich. Pierce C18 tips (Cat #) were purchased from Thermo Scientific.

### NB conjugation

#### Preparation of stock solutions

Peptide solution (4.2 mM): 1mg of angiotensin dissolved in 227 μL ddH_2_O, lumiflavin Solution (0.78 mM): 1 mg lumiflavin dissolved in 5 mL ddH_2_O and 3-methylene-2-norbornanone (0.2 M): 2.052 μL in 84 μL DMSO.

#### Procedure

To a 1 ml vial, peptide (0.00042 mmol, 1 equiv.) in 100 μL ddH_2_O, lumiflavin (0.000042 mmol, 10 mol%) in 53 μL ddH_2_O were added from the stock solutions. To this reaction vial, 3-methylene-2-norbornanone (0.0042 mmol, 10 equiv.) in 21 μL of DMSO, 42 μL cesium formate buffer (pH 3.5, 0.1 M), and 204 μL ddH_2_O were added. The resulting solution was degassed by sparging with nitrogen for 3 min, parafilmed, and irradiated using 2^nd^ generation Analytica blue LED lamps. The sample was irradiated under 110 mW for 6 hours with stirring at 550 rpm and fan cooling. After 6 hours, the sample was removed, diluted with 523 μL of methanol, filtered. To this solution 1 equiv. of p-Toluic acid (in 57 μL of methanol) was added as an internal standard and subjected to LC-MS.

##### Calculation of reaction conversion

Initial optimization with angiotensin was studied using a Waters Acquity UPLC (Empower Software) connected to an Advion Expression CMS mass spectrometer. Method: Solvent A: Acetonitrile, Solvent B: Water, (5% A and 95% B, 0-2 min; 10% A and 90%, 2-3 min; 20% A and 80% B, 3-10 min; 30% A and 70% B, 10-12 min; 95% A and 5% B, 12-16.5 min; 95% A and 5% B, 16.5-18 min; 5% A and 95% B, 18-20 min). For further optimization of Angiotensin at UT-Austin, Liquid chromatography/Mass Spectra were recorded on an Agilent Technologies 6120 Single Quadrupole or 6130 Single Quadrupole interfaced with an Agilent 1200 series liquid chromatography system equipped with a diode-array detector. Resulting spectra were analysed using Agilent LC/MSD ChemStation. All liquid chromatography experiments were run with a 5-95% gradient elution (methanol/water) over 15 minutes. The conversion was calculated by ratio of peak areas of *C*-terminal modified angiotensin and *p*-toluic acid (internal standard) under UV trace for angiotensin photoredox reaction optimization reaction (**SI Table 1-4**) For determination of conversions of peptides with varying *C*-terminal amino acids (**Table 1**), the value was obtained by dividing the peak area of *C*-terminal modified product by addition of peak areas of *C*-terminal modified product and unreacted starting materials under total ion chromatogram (TIC).

##### Protein digestion

100 μg of mass spec-compatible yeast and human K562 cell protein extracts, and bovine serum albumin (get product numbers) were denatured in 2,2,2-trifluoroethanol (TFE) and 5 mM tris(2-carboxyethyl)phosphine (TCEP) at 55 °C for 45 min. Proteins were alkylated in the dark with 5.5 mM iodoacetamide, and then the remaining iodoacetamide was quenched in 100 mM dithiothreitol (DTT). MS-grade GluC, LysargiNase, LysN or AspN was then added to the solution at enzyme:protein ratios of 1:50, 1:50, 1:50, and 1:25 respectively and the digestion reaction was incubated at 37 °C for 24 hours. Samples were then frozen at −20 °C, thawed, and the volume was reduced to 500 μL in a vacuum centrifuge. Samples were then filtered using a 10 kDa Amicon filter and desalted using Pierce C18 tips (Thermo Scientific). Samples were resuspended in 95% water, 5% acetonitrile, 0.1% formic acid prior to mass spectrometry.

##### Mass spectrometry

Peptides were separated on a 75 μM × 25 cm Acclaim PepMap100 C-18 column (Thermo Scientific) using a 5−50% acetonitrile + 0.1% formic acid gradient over 60 or 120 min (for BSA and the lysate samples respectively) and analyzed online by nanoelectrospray-ionization tandem mass spectrometry on an Orbitrap Fusion (Thermo Scientific). Data dependent acquisition was activated with parent ion (MS1) scans collected at high resolution (120,000). Ions with charge +1 were selected for collision-induced dissociation fragmentation spectrum acquisition (MS2) in the ion trap using a Top Speed acquisition time of 3 s. Dynamic exclusion was activated with a 60-s exclusion time for ions selected more than once. MS proteomics data were acquired in the UT Austin Proteomics Facility and have been deposited to the ProteomeXchange Consortium via the PRIDE partner repository with data set identifiers PXD026393.

##### Protein identification

Proteins were identified with Proteome Discoverer 2.3 (Thermo Scientific), searching against the Uniprot human, yeast, or bovine reference proteome. Modifications of the C-terminus were added with the expected NB mass addition of [+78.083 Da] in addition to methionine oxidation [+15.995 Da], *N*-terminal acetylation [+42.011 Da], *N*-terminal methionine loss [-131.04 Da], and *N*-terminal methionine loss with the addition of acetylation [-89.03 Da]. Peptide and protein identifications were thresholded at a 1% false discovery rate.

##### MSFragger analysis

Raw files from mass spectrometry experiments were converted to the mzML format using MSConvert with the peak picking filter selected. The files were split into their respective biological directories and analyzed in 2 separate batches (one for yeast and one for human). Modifications were identified using MSFragger 2.4 with the yeast proteome UP000002311 (accessed 2/3/2021) and the human proteome UP000005640 (accessed 3/25/2020) containing reversed protein sequences as a decoy database. We performed an open search with a 75ppm fragment mass tolerance and the upper and lower precursor mass windows set to 500 and −200, respectively. *N*-terminal acetylation [+42.01060 Da] and methionine sulfoxidation [+15.99490 Da] were considered as variable modifications. Spectra were assigned to peptides and processed using PeptideProphet with default settings. Protein identification was performed using ProteinProphet with default settings. A false discovery rate for all identification steps was set at 1%.

##### Fluorophore labeling of angiotensin

1 mg (~1 μmole) of angiotensin (in 100 μL) was first labeled with 50 eq of NB-PEG4-alkyne along with lumiflavin (10% mol/mol of angiotensin) in 0.1 M cesium formate buffer using the setup as optimized and described earlier. The negative controls were set up with (a) 1 mg angiotensin and 50 eq NB (lacks alkyne handle) and (b) 1 μmole of inert peptide (amide terminal and no internal acidic residue) with sequence AGAGANGSNFGAN-(CO)NH_2_. After the reaction, we adjusted the pH of the solution to 8.5 using 100 μL of HEPES buffer (pH 8.5) and made up the volume to 600 μL with water. We then incubated these peptide solutions with PCA functionalized Synphase lanterns (Mimotopes) for 16h at 37 C to capture the peptides onto the lanterns. We then performed a series of washes of the lanterns with acetonitrile/water (1:1vv), acetonitrile/water with 0.1% formic acid (1:1vv), acetonitrile and dimethylformamide by soaking the lantern for 5 mins in each solvent. To label the internal aspartic acid - in the first step, we performed amide coupling to conjugate azide group on the amino acid side chain using 50 eq 3-Azidopropylamine (Cat#A2738, TCI), 50 eq HCTU and 50 eq DIEA base in 600 μL DMF and incubating at 37C for 2h. Following the washing of lanterns, we added 1 eq of Atto647N-PEG4-DBCO in a 600 μL of 1:1 vv of DMF/water and incubated the lantern for 16h at room temperature. We again washed the lanterns to remove the excess dyes. The labeled peptides were cleaved by reacting the lanterns with 600 μL of TFA cocktail (95% TFA, 2.5% water and 2.5% triisopropylsilane) at room temperature for 2h. After blowing dry N_2_ to remove the TFA and cold ether precipitation, we solubilized the fluorophore labeled peptides with 50 μL acetonitrile/water mixture. We then de-protected the PCA modified N-terminus of the peptide by incubating in a hydrazine (dimethylaminoethylhydrazine dihydrochloride) at 60 °C for 16h.^15^

##### Peptide surface immobilization

For single molecule peptide sequencing, a 40mm German Desag 263 borosilicate glass coverslip (Bioptechs) surface was first cleaned by UV/ozone (Jelight Company) and then functionalized by soaking for 30 minutes in methanol containing 0.01% azidopropyltriethoxysilane (Gelest, Cat # SIA0777.0) and 4mM acetic acid. Weakly attached silane was removed by gentle agitation for 10 minutes in a bath of methanol and a subsequent 10 minutes in water. The coverslip with immobilized peptides were dried under a nitrogen gas stream and baked in a vacuum oven for 20 minutes at 110C.

##### Fluorosequencing

Angiotensin peptides were covalently coupled to the coverslip surface *via* copper catalyzed click chemistry between the alkyne modified C-terminal amino acid residue and the azido silane. A fresh solution of of 2 mM copper sulfate, 1 mM tris(3-hydroxypropyltriazolylmethyl)amine (Sigma, Cat # 762342), 20 mM HEPES (pH 8.0), 5mM sodium ascorbate with fluorescently labelled angiotensin was incubated for 30 minutes at room temperature on the coverslip, washed with water to remove unbound peptides, and dried under a nitrogen gas stream. Single molecule sequencing was performed as described^9^. Fluorosequencing datasets were analyzed using the *SigProc* software tool, available as part of the Plaster package at https://github.com/erisyon/plaster. The raw image files are uploaded to Zenodo (doi: 10.5281/zenodo.4989148)

## ASSOCIATED CONTENT

The supporting information is available free of charge at http://pubs.acs.org.

## ACKNOWLEDGEMENTS

We gratefully acknowledge the UT Mass Spectrometry Facility for their assistance. This work was supported by the Welch Regents Chair (F-0046) to E.V.A. and support from Erisyon, Inc., to E.M.M. and E.V.A. E.M.M. acknowledges additional support from the Welch Foundation (F-1515), NIH (R35 GM122480, R01 HD085901, R01 DK110520) and NSF STTR Grant (#1938726) with Erisyon Inc. Mass spectrometry data collection was supported by CPRIT grant RP110782 to Maria Person. J.S., E.M.M., and E.V.A. are co-founders and shareholders of Erisyon, Inc., and E.M.M. and E.V.A. serve on the scientific advisory board. J.S, E.M.M, E.V.A, B.M.F and L.Z are co-inventors on a patent application relevant to this work.

