## Supplemental Information for "Photoredox-catalyzed decarboxylative *C*-terminal differentiation for bulk and single molecule proteomics"

**Supplementary Figures**

**SF 1: Observation of C-terminal decarboxylated angiotensin.**
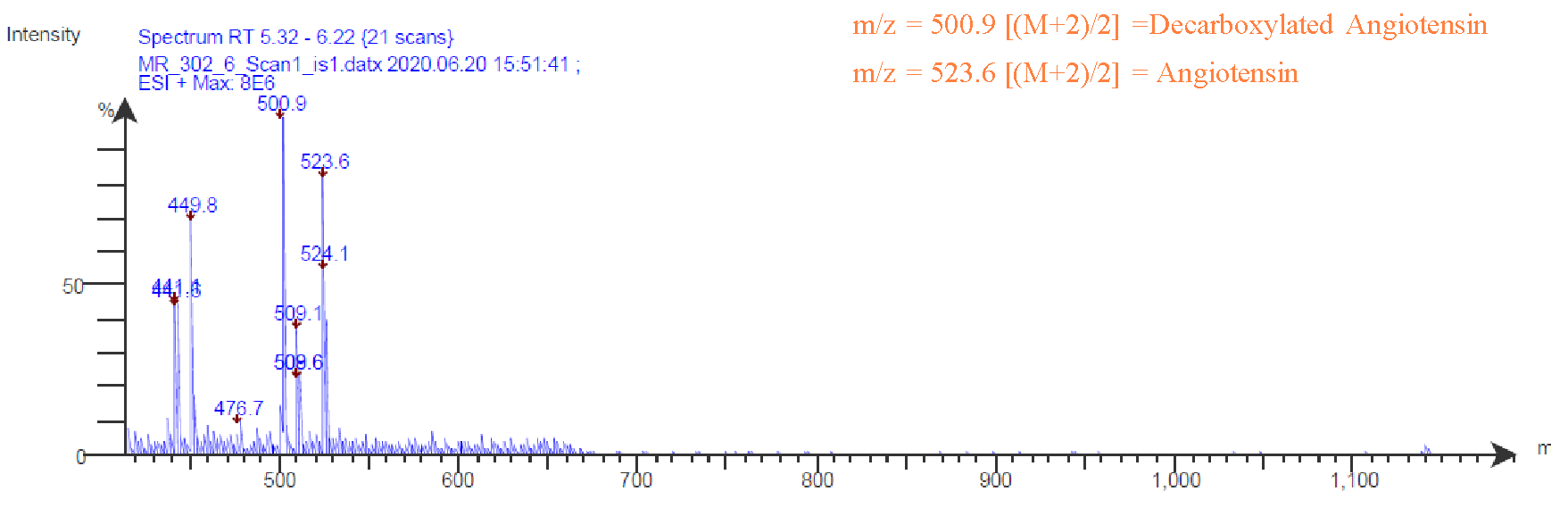

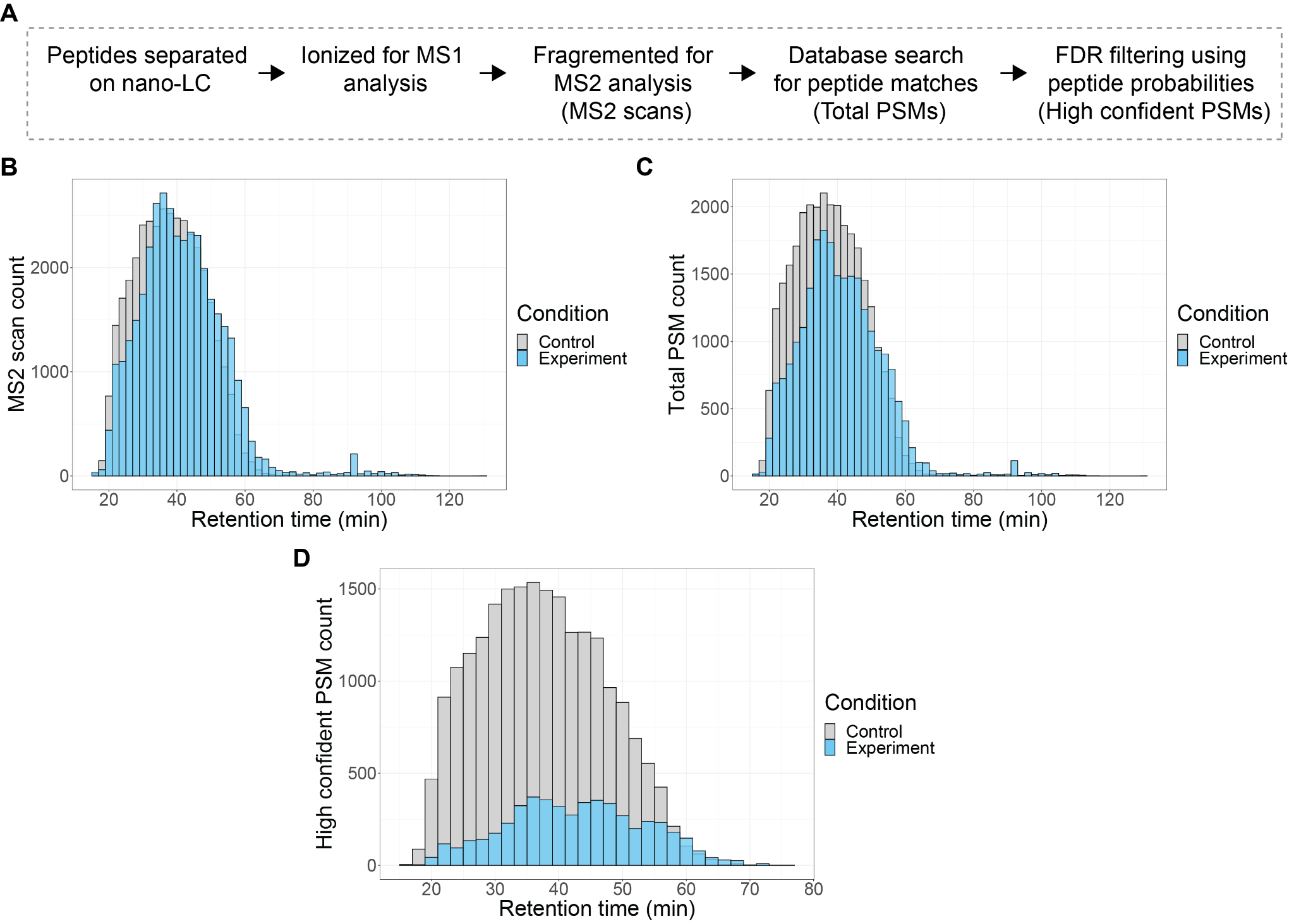

**SF 2: Most of the apparent decrease in identified peptides stems from a decrease in confidence scores for peptide spectral matches (PSMs) in the reference database mass spectral search.** Manual examination of mass spectra suggests that at least part of this effect may stem from the presence of the modification negatively impacting the database search process. In that case, the measured modification efficiencies could thus underestimate the true modification rates. (A) Workflow of the mass spectrometry analysis. Distribution of MS2 scans (B), total PSMs (C), and high confidence PSMs (D) from the mass spectrometry analysis of a trypsin digested yeast cell extract.

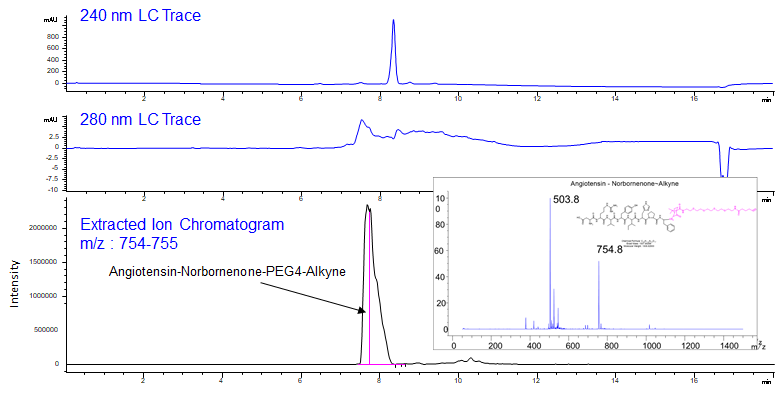

**SF 3: Labeling of angiotensin C-terminus with norbornene-PEG4-alkyne**

**Supplementary Tables**

**Supplementary Table 1 (ST1)**: **Effect of buffer and co-solvent on the reaction**

| **Entry** | **Aqueous buffer** | **Co-solvent** | **% conversion^a^** |
| --- | --- | --- | --- |
| 1 | Cesium formate buffer (pH 3.5) | No co-solvent | 38% |
| 2 | Cesium formate buffer (pH 3.5) | Ethylene glycol (5% v/v) | 24% |
| 3 | Cesium formate buffer (pH 3.5) | *tert*-Butanol (5% v/v) | 37% |
| 4 | Cesium formate buffer (pH 3.5) | Glycerol (5% v/v) | 21% |
| 5 | Cesium formate buffer (pH 3.5) | 1,4-Dioxane (5% v/v) | ND^b^ |
| 6 | Cesium formate buffer (pH 3.5) | ACN (5% v/v) | 43% |
| 7 | Cesium formate buffer (pH 3.5) | DMSO (5% v/v) | 63% |
| 8 | PBS buffer (pH 10.7) | DMSO (5% v/v) | ND^b^ |
| 9 | Phosphate buffer (pH 7.0) | DMSO (5% v/v) | 25% |
| 10 | Sodium acetate buffer (pH 5.1) | DMSO (5% v/v) | 30% |
| 11 | Cesium formate buffer (pH 3.5) | DMSO (2.5% v/v) | 52% |
| 13 | Cesium formate buffer (pH 3.5) | DMSO (10% v/v) | 74% |
| 14 | Cesium formate buffer (pH 3.5) | DMSO (20% v/v) | 70% |
| ^a^ % conversions were based on UPLC data. ^b^ ND : Not determined. | | | |

**Supplementary Table 2 (ST2): Optimization of catalyst loading**

| **Entry** | **Catalyst loading** | **Michael acceptor (equiv.)** | **%conversion**^a^ |
| --- | --- | --- | --- |
| 1 | 1 mol% | 10 | 14% |
| 2 | 3 mol% | 10 | 24% |
| 3 | 5 mol% | 10 | 58% |
| 4 | 7.5 mol% | 10 | 65% |
| **5** | **10 mol%** | **10** | **81%** |
| ^a^ %conversions were based on UPLC data. | | | |

**Supplementary Table 3 (ST3): Stoichiometry of Michael acceptor and concentration of reaction**

| **Entry** | **Concentration** | **Michael acceptor (equiv.)** | **%conversion^a^** |
| --- | --- | --- | --- |
| **1** | 1 mM | 1 | 46% |
| **2** | **1 mM** | **10** | **81%** |
| 3 | 1 mM | 20 | 79% |
| 4 | 0.5 mM | 10 | 80% |
| 5 | 2 mM | 10 | 48% |
| ^a^ % conversions were based on UPLC data. | | | |

**Supplementary Table 4 (ST4): Optimization of reaction time and power**

| **Entry** | **Reaction Time/Power** | **Conversion**^a^ | **Entry** | **Reaction Time/power** | **Conversion**^a^ |
| --- | --- | --- | --- | --- | --- |
| 1 | 3h/110 mW | 34% | 12 | 6h/205 mW | 26% |
| 2 | 4h/110 mW | 37% | 13 | 3h/255 mW | 46% |
| 3 | 5h/110 mW | 44% | 14 | 4h/255 mW | 37% |
| 4 | **6h/110 mW** | **60%** | 15 | 5h/255 mW | 33% |
| 5 | 7h/110 mW | 26% | 16 | 6h/255 mW | 18% |
| 7 | 8h/110 mW | 22% | 17 | 3h/160 mW | 49% |
| 8 | 9h/110 mW | 18% | 18 | 4h/160 mW | 54% |
| 9 | 3h/205 mW | 42% | 19 | 5h/160 mW | 49% |
| 10 | 4h/205 mW | 44% | 20 | 6h/160 mW | 40% |
| 11 | 5h/205 mW | 56% |  |  |  |
| ^a^ Conversions were determined by radio of peak area of product and internal standard under UV trace | | | | | |
